# Deep repertoire mining uncovers ultra-broad coronavirus neutralizing antibodies targeting multiple spike epitopes

**DOI:** 10.1101/2023.03.28.534602

**Authors:** Jonathan Hurtado, Thomas F. Rogers, David B. Jaffe, Bruce A. Adams, Sandhya Bangaru, Elijah Garcia, Tazio Capozzola, Terrence Messmer, Pragati Sharma, Ge Song, Nathan Beutler, Wanting He, Katharina Dueker, Rami Musharrafieh, Michael J.T. Stubbington, Dennis R. Burton, Raiees Andrabi, Andrew B. Ward, Wyatt J. McDonnell, Bryan Briney

## Abstract

Development of vaccines and therapeutics that are broadly effective against known and emergent coronaviruses is an urgent priority. Current strategies for developing pan-coronavirus countermeasures have largely focused on the receptor binding domain (**RBD**) and S2 regions of the coronavirus Spike protein; it has been unclear whether the N-terminal domain (**NTD**) is a viable target for universal vaccines and broadly neutralizing antibodies (**Abs**). Additionally, many RBD-targeting Abs have proven susceptible to viral escape. We screened the circulating B cell repertoires of COVID-19 survivors and vaccinees using multiplexed panels of uniquely barcoded antigens in a high-throughput single cell workflow to isolate over 9,000 SARS-CoV-2-specific monoclonal Abs (**mAbs**), providing an expansive view of the SARS-CoV-2-specific Ab repertoire. We observed many instances of clonal coalescence between individuals, suggesting that Ab responses frequently converge independently on similar genetic solutions. Among the recovered antibodies was TXG-0078, a public neutralizing mAb that binds the NTD supersite region of the coronavirus Spike protein and recognizes a diverse collection of alpha- and beta-coronaviruses. TXG-0078 achieves its exceptional binding breadth while utilizing the same VH1-24 variable gene signature and heavy chain-dominant binding pattern seen in other NTD supersite-specific neutralizing Abs with much narrower specificity. We also report the discovery of CC24.2, a pan-sarbecovirus neutralizing mAb that targets a novel RBD epitope and shows similar neutralization potency against all tested SARS-CoV-2 variants, including BQ.1.1 and XBB.1.5. A cocktail of TXG-0078 and CC24.2 provides protection against *in vivo* challenge with SARS-CoV-2, suggesting potential future use in variant-resistant therapeutic Ab cocktails and as templates for pan-coronavirus vaccine design.

## Introduction

In the past two decades, three novel coronaviruses (**CoVs**) have crossed from zoonotic hosts into humans and acquired the ability to spread by human-to-human transmission. Two of these pandemic CoVs, severe acute respiratory syndrome (**SARS**)-CoV in 2002 and Middle East respiratory syndrome (**MERS**)-CoV first identified in 2012, have caused relatively small outbreaks of human respiratory disease [1,2]. The third, SARS-CoV-2, resulted in a pandemic of CoV disease 2019 (**COVID-19**) with higher global mortality than any acute virus outbreak in over a century [3].

Current vaccines provided protection against the ancestral strain of SARS-CoV-2 and are still effective at preventing hospitalization but provide less protection against newly emerged variants [4]. Furthermore, the ongoing emergence of increasingly diverse variants of concern (**VoCs**) and the possibility of future spillovers of novel CoVs make pan-CoV vaccine development a public health issue of the highest priority. SARS-CoV-2 infection and vaccination both induce antibody (**Ab**) and T cell-mediated immunity that is associated with reduced disease severity and transmission [5,6]. These responses develop in humans as immune ensembles—polyclonal populations of naive and recalled antigen-experienced immune cells which can target new, previously seen, or evolutionarily-related antigens by searching an extremely diverse Ab repertoire space [7–9]. In response, many pathogens have developed sophisticated immune evasion mechanisms that conceal neutralizing epitopes or distract the Ab response with highly immunogenic, non-neutralizing epitope regions [10]. Thus, broad and potently neutralizing Abs against such “evasion-strong” viruses are typically quite rare and their discovery often requires very deep sampling of the pathogen-specific Ab repertoire [11]. Such antibodies are vitally important, however, both as potential clinical treatments and to inform the design of vaccine immunogens that focus the immune response toward conserved regions of viral vulnerability [12].

Most known Abs against SARS-CoV-2 recognize one of three major epitope regions: the receptor-binding domain (**RBD**), the S2 subunit, and the N-terminal domain (**NTD**). Neutralizing Abs (**nAbs**) have been reported against each of these regions [13,14], with the broadest nAbs targeting epitopes in S2 [15–19] or the RBD [20,21]. In contrast, NTD-specific Abs primarily target a single supersite using a limited genetic vocabulary [22,23] and, because the NTD is not particularly well conserved across CoVs, are presumed to be of limited breadth [24]. Recently reported NTD-specific Abs targeting a moderately conserved epitope outside the NTD supersite neutralized several SARS-CoV-2 variants of concern (**VoCs**), but their breadth did not extend beyond SARS-CoV-2 to other human CoVs [25].

Here, we employ a multiplexed antigen screening method using single cell immune profiling technology, which facilitates the rapid recovery of thousands of natively paired Ab sequences together with detailed antigen specificity information (***Fig 1a***). We applied this method to a single convalescent COVID-19 survivor and to a small cohort of longitudinally sampled COVID-19 patients who were infected early in the pandemic and subsequently vaccinated, yielding more than 9,000 SARS-CoV-2-specific monoclonal Abs (**mAbs**). This comprehensive survey enables targeted discovery of extremely rare but desirable Ab specificities while simultaneously providing an expansive view of the pathogen-specific Ab repertoire. The rich single cell profiles generated with this workflow allow fine-grained deconvolution of Ab binding patterns, revealing that increased Ab breadth following vaccination of previously infected individuals is due to the broadening of individual mAbs rather than supplementation by narrow but complementary mAb specificities. We observed substantial clonal convergence among SARS-CoV-2-specific mAbs, in which mAbs from different individuals coalesce upon similar “public” genetic solutions to pathogen recognition. We recovered two particularly interesting nAbs: CC24.2, which targets a novel RBD epitope and exhibits pan-sarbecovirus breadth that includes BA, BQ, and XBB variants of the SARS-CoV-2 Omicron lineage; and TXG-0078, an NTD-specific nAb able to recognize a diverse range of alpha- and beta-CoVs. TXG-0078 alone, as well as a cocktail of TXG-0078 and CC24.2, protects against *in vivo* SARS-CoV-2 challenge when given prophylactically. Broadly protective mAb cocktails are in some ways preferrable to monotherapy, as increased epitope diversity provides added protection against viral escape. The discovery of these novel conserved neutralizing epitopes will also inform the future development of pan-CoV vaccines.

**Figure 1.**
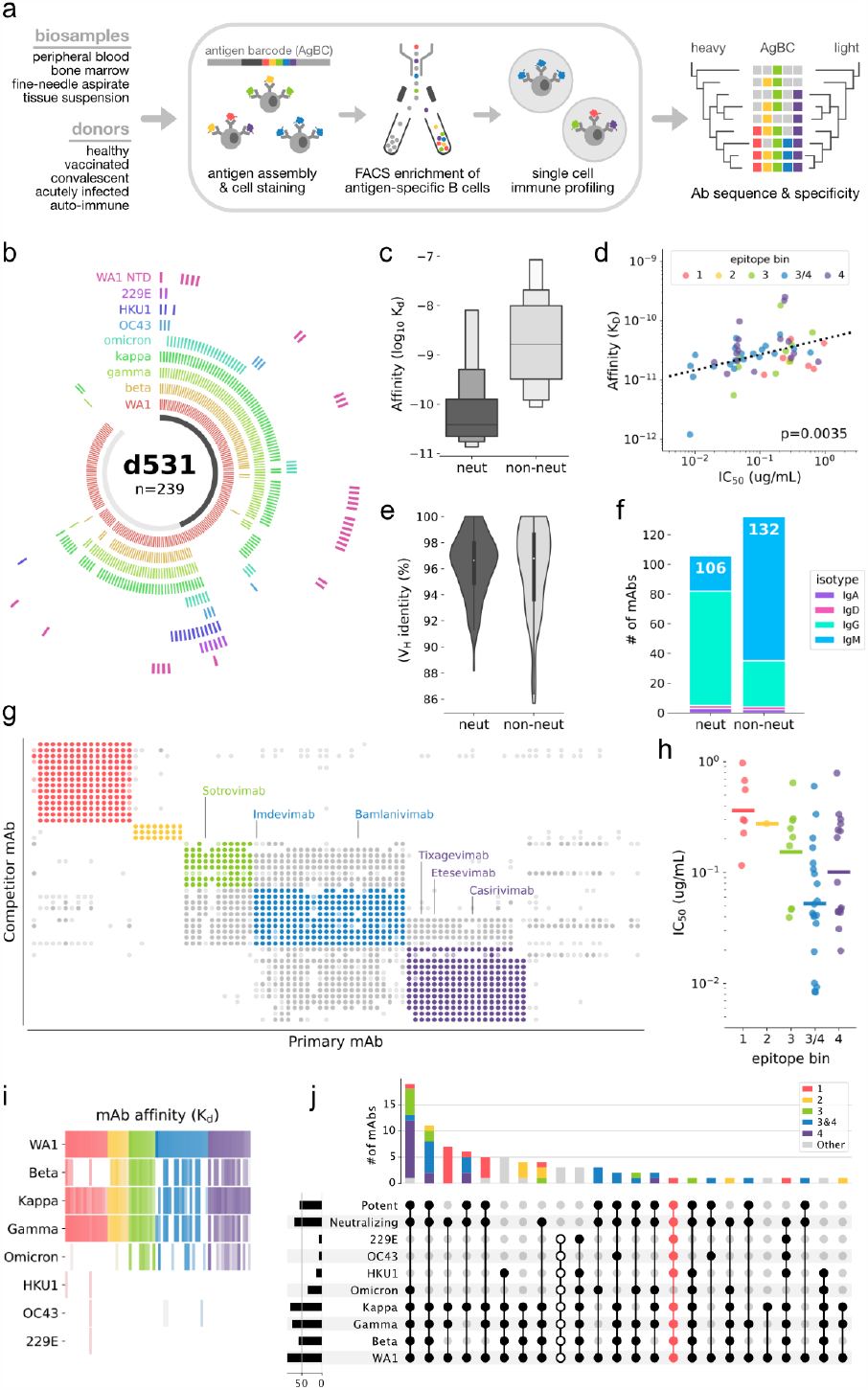
Discovery of an ultra-broad neutralizing antibody targeting the NTD. (a) Overview of the multiplexed antigen screening method. (b) Binding specificities of recombinantly expressed mAbs from Donor 531 (d531). Each arc segment corresponds to a single mAb, and the ring corresponding to each Spike antigen is colored if bound by the mAb. The centermost band is colored dark gray if the mAb neutralizes SARS-CoV-2 and light gray if non-neutralizing. (c) Letter-value plot of mAb affinity for WA1 Spike. The center line on each box represents the median. (d) Correlation between binding affinity (KD) and neutralization (IC50) for the WA1 strain of SARS-CoV-2. The linear regression line is shown, as well as the p-value indicating a significant positive slope. (e) Violin plot of heavy chain Variable (VH) germline gene identity. 100% identity indicates no somatic mutation. (f) Isotype distribution of WA1 nAbs and non-nAbs. The total number of mAbs in each category (*neut* and *non-neut*) is shown at the top of the respective bar. (g) Epitope binning by competitive binding inhibition. For each mAb pair, a darkly filled circle indicates strong competition, a lightly filled circle indicates weak competition, and no circle indicates no competition. Points are colored according to their respective epitope bin. (h) Neutralization potency against SARS-CoV-2 of nAbs in each epitope bin. Bins are colored as in (g). (i) Heatmap of mAb binding affinity (KD) for SARS-CoV-2 VoCs and seasonal CoV S-proteins. Colors correspond to epitope bins and are the same as in (g). Color intensity corresponds to binding affinity, with high affinity colored most intensely. (j) Upset plot showing cross-reactivity binding and neutralization patterns among epitope binned mAbs. The broadest cross-reactivity group, which bind all tested CoV S-proteins but do not neutralize, are indicated by unfilled dots. TXG-0078, which comprises its own cross-reactivity pattern, is highlighted in pink.

## Results

We first obtained approximately 10^8^ peripheral blood mononuclear cells (**PBMCs**) from a convalescent participant with prior PCR-confirmed SARS-CoV-2 infection. Using a small, multiplexed antigen bait panel consisting of soluble prefusion trimeric WA1 Spike (**S-protein**), prefusion trimeric D614G S-protein, and human serum albumin (**HSA**) as a negative control, antigen-specific B cells were enriched by flow cytometric sorting and processed using the 10x Genomics Chromium platform, recovering 2,737 natively paired mAbs. We selected a subset of 239 mAbs for recombinant expression and detailed characterization (***Fig 1b***).

Regardless of the original isotype, all mAb variable regions were recombinantly expressed using a IgG1 backbone vector. Most mAbs detectably bound SARS-CoV-2 (197 of 239; 82%), demonstrating the high purity with which this workflow enriches antigen-specific B cells. It is important to note that this binding fraction likely underestimates the selectivity of this method. Non-binding mAbs were overwhelmingly IgM (34 of 42; 81%) and it is possible that some, perhaps many, of these mAbs rely more heavily on avidity for S-protein binding. Avidity-driven interactions would likely be overlooked when evaluating these mAbs as recombinant IgGs. Binding mAbs were generally of high affinity, with 130 of 197 (66%) mAbs producing apparent KD values in the picomolar range.

Of the 239 recombinantly expressed mAbs, 106 (44%) neutralized the WA1 strain of SARS-CoV-2 in a live virus red dye uptake assay. As a group, neutralizing mAbs (nAbs) bound soluble WA1 Spike with higher affinity than did non-nAbs (***Fig 1c***). Binding affinity correlated with neutralization potency (***Fig 1d***), indicating that low affinity may be at least partially responsible for the lack of detectable neutralization by non-nAbs. Although nAbs and non-nAbs showed similar levels of somatic mutation (***Fig 1e***), nAbs were predominantly IgG (73%) while a similarly sized majority of non-nAbs were IgM (73%; ***Fig 1f***). This parallels the results seen with non-binding mAbs and again suggests that avidity-reliant IgM mAbs may not reproduce their native function when expressed recombinantly as IgGs. Epitope binning by competitive binding inhibition identified five major specificity groups (***Fig 1g***). Three groups (3, 3/4, and 4) represent the RBD epitopes targeted by marketed therapeutic mAbs and the remaining two groups (1 and 2) correspond to distinct epitope regions on the NTD. Neutralizing mAbs against RBD epitopes were generally more potent than NTD nAbs, with the most potent nAbs belonging to competition bin 3/4 (***Fig 1h***).

Screening against a diverse panel of CoV S-proteins, including several SARS-CoV-2 VoCs and seasonal CoVs, revealed a variety of cross-reactivity binding patterns (***Fig 1i-j***). The Beta S-protein escaped recognition by most mAbs in epitope bin 1, and very few NTD mAbs (epitope bins 1 and 2) were able to bind the Omicron S-protein (***Fig 1i***). Although a handful of mAbs were able to recognize seasonal CoVs, binding was much weaker. To assess cross-reactivity, we grouped the mAbs by their ability to neutralize WA1 and bind to a variety of CoV S-proteins. The largest group contained mAbs that neutralize WA1 and bind the S-proteins of all tested SARS-CoV-2 VoCs, including Omicron (***Fig 1j***). This is particularly notable because the donor sample was collected over a year before the Omicron variant was first detected. The only mAbs able to recognize all SARS-CoV-2 VoC and seasonal CoV S-proteins were non-neutralizing, although one especially interesting NTD-specific mAb, named TXG-0078 and highlighted in pink in ***Fig 1j***, bound all tested isolates except Omicron and potently neutralized SARS-CoV-2. TXG-0078 uses the IGHV1-24 germline gene segment, which is characteristic of mAbs targeting the NTD supersite [22,25]. Despite this similarity to existing mAbs, TXG-0078 is the only known neutralizing mAb against the NTD supersite with binding breadth that spans alpha- and beta-CoVs.

### Development of antibody breadth following infection and vaccination

We next assessed the SARS-CoV-2-specific Ab response using a larger multiplexed antigen panel designed to quantify cross-VoC binding breadth. Using a longitudinal cohort of participants who had been infected with SARS-CoV-2 (post-infection timepoints T1, T2 and T3) and subsequently vaccinated (post-vaccination timepoint T4), we isolated a total of 6,302 paired mAbs (***Fig 2a***). The isotype distribution was similar across all four participants, and although the majority of SARS-CoV-2-specific mAbs were IgM (58%, ***Fig 2b***), the use of IgG increased markedly with breadth (18.3% in single-variant binders, 65.6% in pan-VoC binders; ***Fig 2c***). Heavy chain variable gene (**V-gene**) use among SARS-CoV-2-specific mAbs was quite diverse, and we did not observe substantial enrichment of specific V-genes among cross-reactive mAbs (***Fig 2d***). Broadly reactive mAbs were more somatically mutated than single-variant binders (***Fig 2e***), which, together with the isotype composition observations, suggests an association between increased maturation (class-switching and somatic hypermutation) and binding breadth. Interestingly, although somatic mutation frequency increased consistently over time (***Fig 2f***), mAb breadth remained relatively static following infection before substantially increasing after vaccination for three of four participants (***Fig 2g***).

**Figure 2.**
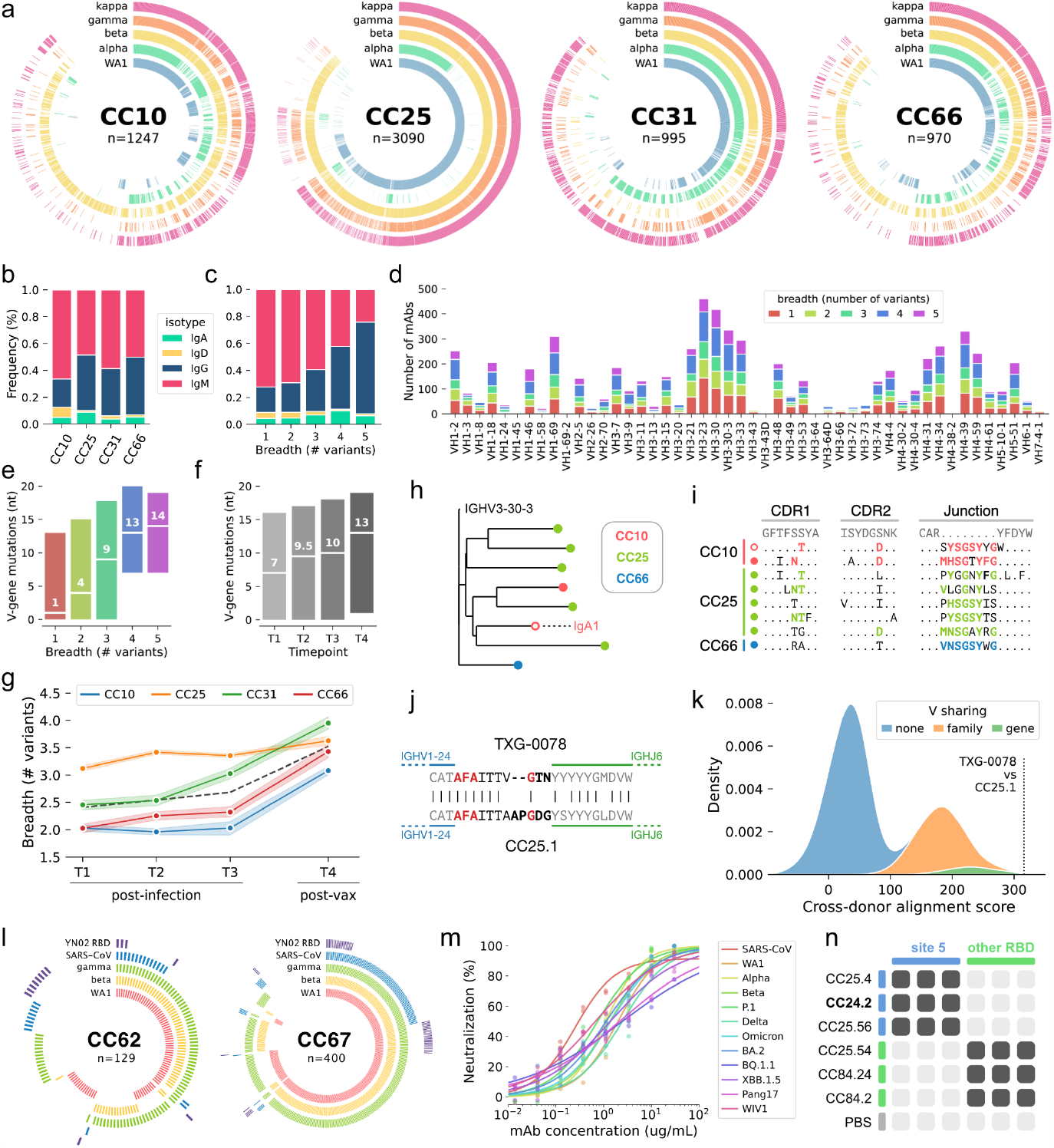
Quantifying the development of mAb breadth of binding using antigen barcodes. (a) Ring plots showing the antigen barcode classifications of 6,305 SARS-CoV-2-specific mAbs from participants CC10, CC25, CC31 and CC66. Individual mAbs are represented by radial vectors, with each antigen ring (WA1: blue, alpha: green, beta: yellow, gamma: orange, kappa: magenta) colored if the given B cell was classified as positive for the respective antigen. Antibodies are sorted by breadth, with breadth decreasing in the clockwise direction. (b) Isotype distribution of SARS-CoV-2-specific mAbs for each donor. (c) Isotype distribution of mAbs, grouped by cross-variant breadth. (d) Heavy chain Variable (VH) gene use of SARS-CoV-2-speicific mAbs, grouped by cross-variant breadth. (e) Number of VH nucleotide mutations in SARS-CoV-2-specific mAbs, grouped by breadth. (f) Number of VH nucleotide mutations in SARS-CoV-2-specific mAbs, grouped by timepoint (T1, T2 and T3 are post-infection but pre-vaccination; T4 is post-infection and post-vaccination). (g) Median cross-variant breadth of all mAbs isolated at each timepoint, grouped by donor. The dashed black line represents the median of all mAbs from participants. (h) Phylogenetic representation of a single public SARS-CoV-2-specific clonotype. All mAbs in the clonotype use the IgG1 isotype except for a single IgA1 clone (indicated by text and an unfilled marker). (i) Complementarity determining region (CDR) alignments of sequences in the public clonotype shown in (h). Mutations (CDR1 and 2) or untemplated residues (Junction) shared between multiple participants are highlighted in color (CC10: magenta, CC25: green, CC66: blue). The lone IgA1 mAb is indicated by an unfilled marker. (j) Junction alignments of a second public clonotype, comprising TXG-0078 (from donor 531) and CC25.1 (from donor CC25). Untemplated positions (N- and P-addition) are indicated in bold, and identical residues in untemplated regions are highlighted in red. (k) Distribution of cross-donor alignment scores obtained by iterative pairwise alignment of heavy chain amino acid sequences. Only sequences recovered from different doors were compared. Distributions resulting from comparison of sequences encoding the same V gene (green), V genes from the same family (orange) or V genes from different families (blue) are shown. The alignment score between TXG-0078 and CC25.1 is highlighted. (l) Ring plots showing antigen barcode classifications of two additional participants (CC62 and CC67). Antigen rings are colored if the respective mAb was classified as antigen positive (WA1: magenta, beta: yellow, kappa: green, SARS-CoV: blue, YN02 RBD: purple). Neutralization of SARS-CoV (red) and several SARS-CoV-2 variants by mAb CC24.2. (n) Epitope binning of mAb CC24.2 with several RBD-specific mAbs. Two competitor mAbs (CC25.4 and CC25.56) bind the Site 5 epitope, the other competitor mAbs (CC25.54, CC84.24 and CC84.2) bind other RBD epitopes. Binding inhibition (dark gray) or lack of binding inhibition (light gray) is shown, with phosphate buffered saline (PBS) used as a negative control.

### Frequent clonal coalescence among SARS-CoV-2 antibodies

To quantify the occurrence of “public” CoV-specific mAbs, which are highly similar mAbs present in multiple different individuals, we performed pairwise comparisons of every possible combination of CoV-specific mAbs from different participants and ranked them by similarity. Sequences from one such group of public mAbs, isolated from three different participants, showed maturation trajectories of sufficient similarity that cross-donor sequences were phylogenetically interspersed (***Fig 2h***). Indeed, the similarity between these public Ab sequences was not attributable solely to the use of identical germline gene segments but was also the result of substantial homology in non-germline encoded residues (***Fig 2i***). We refer to this phenomenon – sequences with similar germline rearrangements which also follow similar maturation trajectories – as clonal coalescence. We also observed clonal coalescence between TXG-0078, the exceptionally broad NTD-specific nAb described above, and another CoV-specific mAb, CC25.1, which was isolated from the post-vaccination sample of donor CC25 (***Fig 2a, 2j***). Like the coalescence group shown in ***Fig 2h-i***, the heavy chain junctions of TXG-0078 and CC25.1 were remarkably similar even at non-templated positions, resulting in much greater similarity than would be expected by merely encoding the same germline components (***Fig 2k***). TXG-0078 and CC25.1 were isolated at different facilities which did not share samples or reagents, eliminating the possibility of contamination as a cause of the observed clonal coalescence.

### Isolation of a pan-sarbecovirus RBD nAb using an expanded antigen panel

Encouraged by the increased breadth observed in participants who had been both infected and vaccinated, we subsequently screened additional participants using a modified antigen panel designed to assess breadth of binding beyond SARS-CoV-2 variants. This panel included S-proteins from the WA1, beta and gamma strains of SARS-CoV-2, as well as SARS-CoV S-protein and the RBD from YN02, a clade 2 bat coronavirus closely related to SARS-CoV-2 (***Fig 2l***). We observed a surprisingly high frequency of mAbs that bound both SARS-CoV-2 and SARS-CoV (10-20% of all mAbs). One particularly broad mAb, named CC24.2, neutralized SARS-CoV and a broad range of SARS-CoV-2 variants, including Omicron, BA.2, BQ.1.1 and XBB.1.5 (***Fig 2m***). The similar potency with which CC24.2 neutralized all tested sarbecovirus isolates suggests either high conservation of critical contact residues or sufficient binding flexibility that some epitope variability can be tolerated. We also note the near perfect functional complementarity between TXG-0078 and CC24.2: TXG-0078 binds a broad range of CoVs but cannot recognize SARS-CoV or Omicron variants of SARS-CoV-2, while CC24.2 does not exhibit breadth beyond sarbecoviruses but neutralizes SARS-CoV and all tested Omicron variants. Epitope mapping of CC24.2 by competitive binding indicates a specificity that overlaps two mAbs against a novel RBD epitope (reported in a companion manuscript) which has been termed Site 5 (***Fig 2n***).

### Structural definition of a broadly conserved NTD neutralizing epitope

Although many NTD-specific nAbs have been reported, none have demonstrated the binding breadth of TXG-0078, making the specific molecular interactions between TXG-0078 and S-Protein of particular interest. Our cryo-EM analysis of TXG-0078 in complex with the WA1 SARS-CoV-2 Spike resulted in 2 different reconstructions of the spike-Fab complex (3.14 Å and 3.34 Å), displaying the conformational flexibility of the Fab-NTD region (***Fig 3a-b***). We further performed local refinement in the NTD-Fab region, which yielded a 3.9Å reconstruction (***Fig 3c***). TXG-0078 binds the N3 and N5 loops of the NTD using all three heavy chain complementarity determining region (**HCDR**) loops. Several aspects of TXG-0078 binding are similar to 4A8, a previously reported NTD-specific nAb [14]. Both mAbs use a similar angle of approach, encode the same heavy chain V-gene, and share binding interactions between NTD and their HCDR1 and HCDR2 loops. The shared interactions include R246 contacts with HCDR1 residues Y27 and E31, H146 H-bond with T30 on HCDR1, salt-bridge formation between K147 and residue E72 on HCDR2 and K152 H-bond with Y111 on HCDR3. However, the HCDR3 of the two mAbs differ at multiple residues allowing TXG-0078 to make additional contacts not observed with 4A8. Notably, TXG-0078 has a phenylalanine at position 100 on the HCDR3 instead of a threonine in 4A8, resulting in CH/*π* interactions with P251 on the NTD while also forming stabilizing contacts with Y27 on HCDR1. It is possible that these additional HCDR3 interactions allow TXG-0078 to accommodate variability at other contact positions in a way that less broad NTD-specific mAbs cannot.

**Figure 3.**
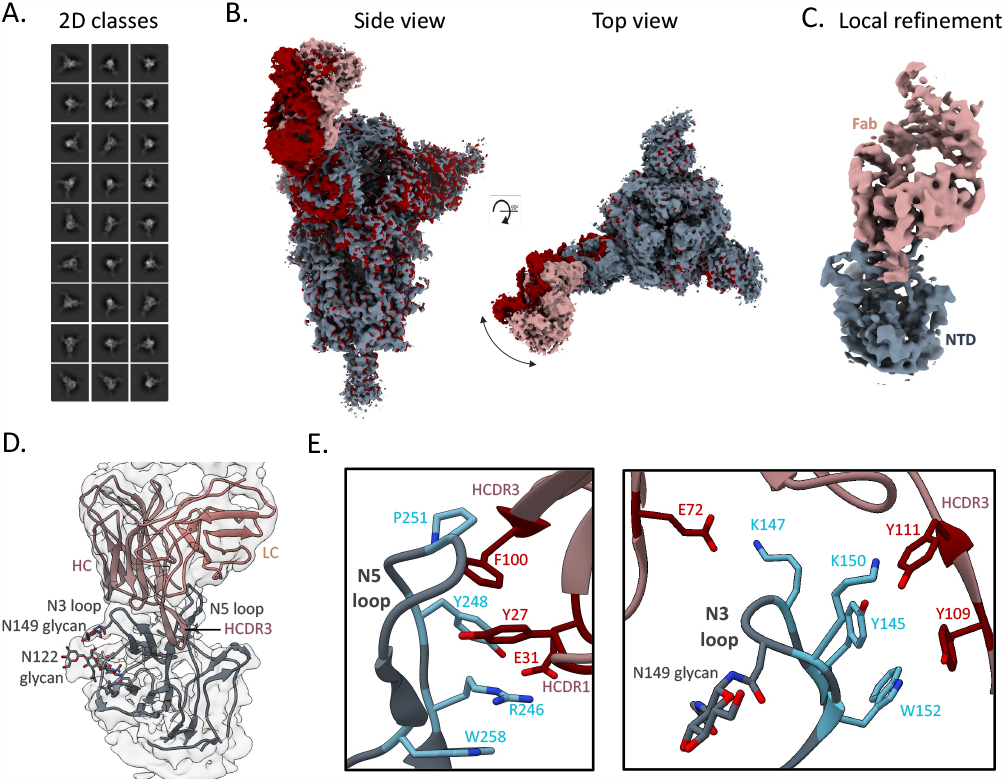
Structural definition of the broad, NTD-supersite antibody TXG-0078. (a) Representative 2D classes obtained by cryo-EM analysis of TXG-0078 Fab in complex with SARS-CoV-2 spike (b) Two cryo-EM reconstructions of the TXG-0078-spike complex at 3.14 Å (Rosy Brown/Grey) and 3.34 Å (Red) resolutions showing conformational flexibility of the Fab-NTD region (c) Cryo-EM reconstruction (3.9Å) of the NTD-Fab region obtained by local refinement. The spike NTD is colored grey and the Fab is colored rosy brown (d) The atomic model of TXG-0078 Fab Fv bound to NTD docked in the local refinement map. (e) The atomic model reveals that TXG-0078 targets the NTD supersite as has been described for other antibodies, though it is capable of weakly binding endemic human coronaviruses. The N3 and N5 loops comprise the epitope of TXG-0078; the CDRH3 of TXG-0078 reaches into a pocket in the N5 loop and the light chain is only minimally engaged in binding. Epitope and paratope residues are colored in light blue and red, respectively.

### Antibody-mediated protection against SARS-CoV-2 infection

As described above, TXG-0078 neutralizes SARS-CoV-2 and binds a diverse array of alpha- and beta-CoVs including OC43, HKU1 and 229E; however, TXG-0078 was unable to neutralize SARS-CoV or Omicron variants of SARS-CoV-2. CC24.2 neutralizes all tested sarbecoviruses, including SARS-CoV and Omicron variants of SARS-CoV-2. Due to their complementary specificities, we evaluated the potential of TXG-0078 and CC24.2 to protect against *in vivo* SARS-Cov-2 challenge alone or in combination. We administered 300μg of prophylactic mAb to a total of 20 ACE2 transgenic mice in four treatment groups: TXG-0078, CC24.2, a two-mAb cocktail comprising 150μg each of TXG-0078 and CC24.2, and ZIKV-1, a negative control antibody targeting Zika virus. Following mAb administration, mice were challenged with the WA1 strain of SARS-CoV-2 through intranasal administration. Each mouse was weighed at day 0 (baseline) and again daily for 5 days before sacrifice. All mice receiving TXG-0078 or the cocktail were protected from weight loss (***Fig 4***). CC24.2 appeared to provide partial protection, with 3 of 5 mice exhibiting a reduction in weight loss comparable to animals receiving TXG-0078 or the cocktail. The reduced efficacy of CC24.2 can likely be attributed to its approximately 10-fold lower potency against SARS-CoV-2 compared to TXG-0078 (IC50 values of 1.63ug/mL and 171 ng/mL, respectively).

**Figure 4.**
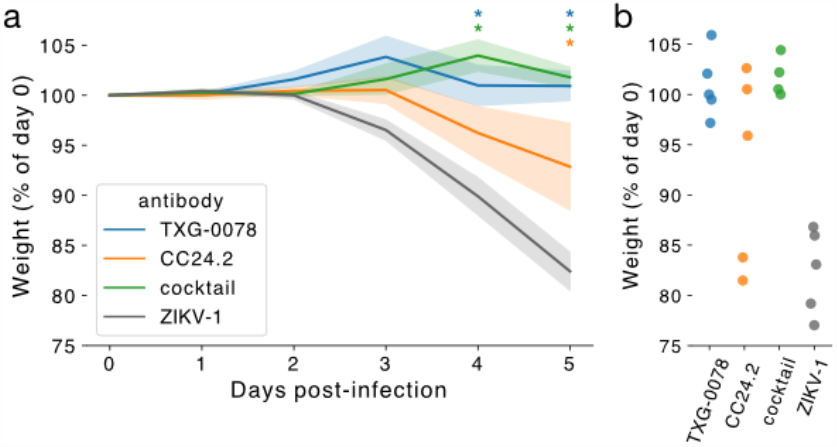
TXG-0078 protects transgenic ACE2 mice from live SARS-CoV-2 challenge. (a) Mice were administered 300 μg of TXG-0078, CC24.2, a cocktail of both TXG-0078 and CC24.2, or a control antibody targeting Zika virus (ZIKV-1). Mean percent change in baseline weight is shown for each group. Shading indicates standard error. Significant differences between groups (p≤ 0.05) are indicated by asterisks in the color of each group displaying significantly different weight loss than the control (ZIKV-1) group. Significance was determined using a one-way ANalysis Of VAriance (ANOVA) with Dunnet’s multiple comparisons test. (b) Percent of baseline weight at day 5 is shown for all mice, separated by treatment group.

## Discussion

As the COVID-19 pandemic made painfully clear, the development of broad, variant-resistant vaccines and therapeutics is urgently necessary to prevent and control future outbreaks of infectious diseases with pandemic potential. Integral to rational vaccine and therapeutics discovery strategies is the identification of conserved regions of viral vulnerability. Unfortunately, conserved neutralizing epitopes are often cryptic or poorly immunogenic and potent Abs targeting such regions can be extremely rare. Here, we demonstrate the power of multiplexed antigen screening using single cell immune profiling technology for deeply surveying and cataloging functional adaptive immune repertoires, by isolating over 9,000 CoV-specific human mAbs. This expansive mAb dataset revealed multiple instances of clonal coalescence, in which different participants independently produced antigen-specific mAbs with remarkable genetic similarity. Of particular interest was TXG-0078, a public NTD-specific neutralizing mAb that binds a broad range of alpha- and beta-coronaviruses and which provided protection against *in vivo* challenge with SARS-CoV-2. The discovery of a conserved neutralizing NTD epitope has exciting implications for the development of therapeutic mAb cocktails that improve resistance to viral escape by targeting multiple conserved epitopes as well as the development of vaccines that protect against current, emerging, and future human CoVs. We also report the discovery of CC24.2, a neutralizing mAb that achieves pan-sarbecovirus breadth by targeting a novel RBD epitope.

Beyond SARS-CoV-2, the ability to perform large-scale functional analyses of circulating B cells allows much more sophisticated and granular analysis of the pathogen-specific Ab response than can be achieved using traditional serology or mAb isolation techniques. With serology, for example, it is not possible to determine whether broadly reactive serum is the result of broadly reactive mAbs or a complementary combination of narrow specificities. Using this single cell multiplexed antigen screening approach, we can interrogate the genetics and binding specificities of thousands of B cells simultaneously, producing an expansive representation of the pathogen-specific humoral response at single mAb resolution. This approach has the potential to greatly advance our understanding of infectious disease biology; speed the discovery of exceptionally rare mAbs with unique, therapeutically desirable specificities; and facilitate the design and evaluation of vaccines that elicit durable, variant-resistant immunity.

## Acknowledgements

The authors would like to thank all the COVID-19 survivors and vaccines who graciously provided the samples used in this study.

## Funding

This work was funded by the National Institutes of Health awards R35-GM133682 (BB), U19-AI135995 (BB), R01-AI170928 (RA), and UM1-AI144462 (DRB). Experiments described in this study were partially funded by 10x Genomics, Inc.

### Author contributions

Conceptualization: W.J.M., B.B., T.R.

Data curation: W.J.M., J.H., S.B., D.B.J., B.A.A, B.B., T.R., A.W.

Formal analysis: W.J.M., D.B.J., S.B., B.B.

Funding acquisition: W.J.M., A.W., R.A., B.B., T.R. Investigation: All authors.

Project administration: W.J.M., T.R., B.B.

Resources: 10x Genomics, Scripps Research Institute, T.R., A.W., B.B. Software: D.B.J., W.J.M., B.B.

Supervision: W.J.M., T.R., B.B.

Validation: all authors. Visualization: W.J.M., B.B., S.B.

Writing – original draft: W.J.M., J.H., T.R., B.B.

Writing – reviewing & editing: all authors.

### Data and materials availability

All source data are available as part of this publication. Ab sequences will be available upon publication via GenBank (Submission #2633676, *accession number is pending*) and via CoV-AbDab. Reagent requests for vectors, other reagents, and data related to these antibodies not made available in this paper may be fulfilled by the corresponding authors on request; requests may require material transfer agreement. Code to replicate key findings of this publication will be available via Github (github.com/briney/10x-CoV-paper).

## Methods

### Recombinant antibody vector synthesis, transient expression, and purification

Variable heavy chain and light chain domains of anti-SARS-CoV-2 S1 antibodies were reformatted to IgG1 and synthesized and cloned into mammalian expression vector pTwist CMV BG WPRE Neo utilizing the Twist Bioscience eCommerce portal. Light chain variable domains were reformatted into kappa and lambda frameworks accordingly. Clonal genes were delivered as purified plasmid DNA ready for transient transfection in HEK Expi293 cells (Thermo Scientific). Cultures in a volume of 1.2 ml were grown for four days, harvested and purified using Protein A resin (PhyNexus) on the Hamilton Microlab STAR platform into 43 mM Citrate 148 mM HEPES, pH 6.

### Carterra LSA surface plasmon resonance experiments

Surface plasmon resonance experiments were performed on a Carterra LSA SPR biosensor equipped with a HC30M chip at 25°C in HBS-TE (10 mM HEPES pH 7.4, 150 mM NaCl, 3 mM EDTA, 0.05% Tween-20). Antibodies were diluted to 10 ug/ml and amine-coupled to the sensor chip by EDC/NHS activation, followed by ethanolamine HCl quenching. Increasing concentrations of ligand were flowed over the sensor chip in HBSTE with 0.5 mg/ml BSA with 5-minute association and 15-minute dissociation. Following each injection cycle the surface was regenerated with 2x 30 second injections of IgG elution buffer (Thermo). Data were analyzed in Carterra’s Kinetics Tool software with 1:1 binding model. Pre-fusion trimeric spike proteins used in the screen included SARS-CoV-2 variants WA-1/2020, beta, gamma, kappa, and omicron, and endemic coronaviruses HKU1, OC-43, and 229E. All antigens were sourced from Acro Biosystems. These reagents contain His tag and T4 foldon domains for purification and trimerization respectively.

### Vero E6 neutralization assays

Assays were performed with a clinical isolate of the SARS-CoV-2 B.1 lineage (MEX-BC2/2020). This virus carries the D614G mutation in the spike protein (full sequence available at the GISAID/EpiCoV database ID: EPI_ISL_747242). The screen was performed with a microneutralization assay that utilizes prevention of the virus-induced cytopathic effect (CPE) in Vero E6 cells. All antibodies were stored at 4°C until its use. The screen was performed in ten different experiments performed over ten days, each one assessing the activity of approximately 30 antibodies in parallel. All plates included a positive control–plasma from a convalescent patient who had also received the first dose of the Pfizer/BioNTech mRNA vaccine (BNT162b2). Plasma was collected 21 days after the vaccine injection. We utilized Vero E6 cells to evaluate neutralization activity against a replication competent SARS-CoV-2 virus. Antibodies were pre-incubated first with the virus for 1h at 37°C before addition to cells. Following pre-incubation of Ab/virus samples, Vero E6 cells were challenged with the mixture. After addition to cells, antibodies were present in the cell culture for the duration of the infection (96 hours), at which time a neutral red uptake assay was performed to determine the extent of the virus-induced CPE. Prevention of the CPE was used as a surrogate marker to determine the neutralization activity of the test-items against SARS-CoV-2. Eight dilutions of each antibody were tested in duplicates for the neutralization assay using a five-fold dilution scheme starting at 1,000ng/mL. When possible, IC50 values of the antibodies displaying neutralizing activity were determined using GraphPad Prism software. Vaccinated plasma control was assessed on each plate using singlet data-points (8 two-fold dilutions throughout 1:20480).

### Cryo-EM analysis

The design and expression of SARS-CoV-2 HP-GSAS Mut7 spike used for cryo-EM studies was done as previously described [26]. The spike protein was incubated with 3-fold molar excess of TXG-0078 Fab (final concentration of 1mg/ml) for 30 minutes and mixed with 0.5 μl of 0.04 mM Lauryl maltose neopentyl glycol (LMNG) solution immediately prior to sample deposition onto plasma-cleaned Quantifoil 1.2/1.3 grids. Grids were then blotted for 4 seconds before being plunged into liquid ethane using a Vitrobot mark IV (Thermo Fisher Scientific). Data collection was performed on a Thermo Fisher Glacios operating at 200 keV mounted with a Thermo Fisher Falcon 4 direct electron detector using the Thermo Fisher EPU 2 software. A total of 4250 micrographs were collected at a total dose of 50 e-/Å2. CryoSPARC Live Patch Motion Correction was used for alignment and dose weighting of movies. CTF estimations, particle picking, particle extraction and iterative rounds of 2D classification were performed on CryoSPARC [27]. The particles were transferred to Relion 3.1 for further processing [28]. Focused classification involving iterative rounds of alignment-free 3D classification with masks around the NTD-Fab region and 3D refinement was used to obtain two reconstructions of the spike-Fab complex. The local refinement of the NTD-Fab region was performed on cryoSPARC with the local refinement job.

### Model building and refinement

Initial model building into the local refinement reconstruction was performed manually in Coot using PDB 7C2L (spike complexed with 4A8 Fab) as a template [29]. Rosetta relaxed refinement was performed on the initial model and EMRinger and MolProbity metrics were calculated following Rosetta refinement to evaluate and identify the best refined models and Phenix comprehensive validation was performed on the final model (28–30).

### Animal challenge study

Each group (n=5) was passively immunized with 300μg of each mAb through intraperitoneal injections 24 hours prior to infection. Mice were infected with 30,000 FFU of SARS-CoV-2 (WA1/2020). Mice were weighed daily, and percent baseline weight was calculated relative to day 0 weight. On day 5, mice were euthanized and lung lobes were extracted for viral titer quantification.

### Single cell multiplexed antigen screening workflow

Approximately 4.4 million enriched B cells (selected magnetically from 100 million PBMCs) were resuspended in labeling buffer (1% BSA in PBS) and underwent Fc blocking for 10 minutes on ice using Human TruStain FcX (BioLegend). Next, cells were stained with the following cocktail of antibodies, antigens and dyes: CD19 PE-Cy7 (clone SJ25C1, BD Pharmingen) for discrimination of CD19+ cells by using fluorescence-activated cell sorting.

Barcoded Antibodies for 10x Single Cell Immune Profiling, which included the following TotalSeq-C oligo barcoded antibodies TotalSeq-C0389 anti-human CD38, TotalSeq-C0154 anti-human CD27, TotalSeq-C0189 anti-human CD24, TotalSeq-C0384 anti-human IgD, TotalSeq-C0100 anti-human CD20, TotalSeq-C0050 anti-human CD19 (clone HIB19, to distinguish it from the flow clone), TotalSeq-C0049 anti-human CD3E, TotalSeq-C0045 anti-human CD4, TotalSeq-C0046 anti-human CD8A, TotalSeq-C0051 anti-human CD14, TotalSeq-C0083 anti-human CD16, TotalSeq-C0090 mouse IgG1 K isotype control, TotalSeq-C0091 mouse IgG2a K isotype control, TotalSeq-C0092 mouse IgG2b K isotype control were used for surface protein profiling.

The final conjugated antigens include TotalSeq-C0951 PE trimerized S (SARS-2), TotalSeq-C0952 PE Human Serum Albumin, TotalSeq-C0956 APC trimerized S D614G (SARS-2), TotalSeq-C0957 APC Human Serum Albumin, and 7AAD for live/dead cell discrimination. Cells were stained in labeling buffer (1% BSA in PBS) in the dark for 30 minutes on ice, then cells were washed 3 times with 2 mL of cold labeling buffer at 350*g for 5 minutes at 4oC, resuspended in cold labeling buffer and a 1:200 addition of live/dead cell discriminating agent 7AAD for 10 minutes on ice in the dark, then washed one more time with labeling buffer at 350*g for 5 minutes at 4oC, then resuspended in labeling buffer and loaded into a Sony MA900 Cell Sorter using a 70 uM sorting chip. Cells were initially gated on being single, live (7AAD-negative) and PE-Cy7-CD19+ and then sorted on their PE and/or APC status directly into master mixed and water based on one of four criteria:

1.**PE+**, representing trimerized S (SARS-2) antigen and/or HSA-binding memory B cells

2.**APC+**, representing trimerized S D614G (SARS-2) antigen and/or HSA-binding memory B cells

3.**PE+ and APC+**, representing a combination of SARS-2, D614G and/or HAS-binding memory B cells

4.**PE and APC negative**, representing cells not binding either SARS-2 antigen or HSA

The resulting volume was adjusted with additional water to match the recommended volume and target for loading with the 10× 5’V2 Single Cell Immune Profiling kit. Flow cytometry data were analyzed using FlowJo. Standard gene expression, V(D)J, and barcoded antigen libraries were constructed using the 10× 5’V2 Single Cell Immune Profiling kit per manufacturer’s instructions. Additional information in this regard can be found at https://www.10xgenomics.com/products/single-cell-immune-profiling. The libraries resulting from the experiments were sequenced on a NovaSeq 6000 using a NovaSeq S4 200 cycles 2020 v1.5 kit, targeting using read 28, 10, 10, and 90 cycles targeting 20,000, 30,000, or 6000 reads per cell for gene expression, barcoded antigen, or Ig libraries, respectively. Libraries were analyzed using cellranger multi per manufacturer recommendation. Sequence analysis identified a total of 239 antibodies based on target and non-target (control) antigen UMI counts.

### Sequencing analysis

Sequencing data were analyzed and visualized using Cell Ranger, enclone, scab, and custom Python and R scripts. Scab is open source and freely available at github.com/briney/scab.

